# An orally administered peptide hydrogel disentangles immune-microbiota crosstalk for long-term ulcerative colitis therapy

**DOI:** 10.64898/2026.08.22.746417

**Authors:** Tianyu Li, Mengqian Shi, Jintao Shen, Peng Zhou, Yaxuan Chen, Lu Yu, Juping Sun, Hao Tang, Qingyang Zhou, Yimeng Du, Bei Tan, Xiaojie Xu, Ruirui Xing, Xuehai Yan

## Abstract

Ulcerative colitis (UC) is a global health challenge driven by immune dysregulation and gut microbiota imbalance.^1^ Current treatments, limited by insufficient efficacy and systemic toxicity during prolonged use, fail to resolve the intertwined immune-microbial pathology.^2^ Here, we report an orally administered self-assembled hydrogel C₁₂-(IIRR)₂I-NH₂ (CIR), engineered from host defense peptides, which disrupts the immune-microbiota entanglement. The CIR hydrogel exhibits structural transformation at the inflamed sites rich in liposaccharide (LPS), a pro-inflammatory molecule derived from pathogenic bacteria. Stable β-sheet nanofibers can transfer to bioactive α-helix conformations, enabling localized therapeutic action with minimal off-target toxicity. In murine colitis models, CIR restores mucosal integrity and suppresses disease severity, outperforming the first-line drug 5-aminosalicylic acid (5-ASA). Microbiome profiling reveals its capacity to rebalance gut microbiota, depleting LPS produced pathogenic bacteria like *Prevotellaceae*. Transcriptomic analyses further indicate that CIR silences TLR4-mediated signaling pathway. By synergistically targeting immune dysregulation and microbial dysbiosis, this self-assembled peptide hydrogel establishes a paradigm-shifting strategy for UC, offering clinically translatable potential for multifactorial gastrointestinal disorders.

## Main

Ulcerative colitis (UC), a chronic inflammatory bowel disease marked by relapsing mucosal inflammation, causes an increasing global health burden due to its rising incidence, lifelong morbidity, and limited therapeutic options.^3,4^ While its pathogenesis involves a complex interplay between immune hyperactivation and gut microbiota dysbiosis, current therapies including aminosalicylates, corticosteroids, and biologics fail to break this vicious cycle.^5–7^ Most of these agents target single pathways (e.g., TNF-α, integrin), leading to decreased efficacy or severe adverse effects in long-term use.^8–12^ Although 5-aminosalicylic acid (5-ASA) is almost the only choice for clinical maintenance treatment, its mechanism of action remains uncertain and the remission rates do not surpass 50% under the shadow of relapsing.^13,14^ As the soil remains dysregulated, relapsing cannot be finely controlled. Thus, an urgent need exists for treatments that resolve the interplay between immune dysregulation and microbial imbalance, while enabling safe, long-term use.

Host defense peptides (HDPs) are natural-derived biomolecules with dual antimicrobial and immunomodulatory functions, which offer a promising blueprint for such therapeutics.^15–17^ Various types of HDPs have been discovered to regulate microbiota composition, dampen excessive inflammation, or enhance barrier integrity, ideally suited for UC treatment.^18,19^ However, their clinical translation has been hindered by protease susceptibility, cytotoxicity, and costly synthesis strategies (e.g., non-natural amino acids).^20,21^

Herein, we present a rationally engineered peptide C12-(IIRR)₂I-NH₂ (CIR), to overcome these barriers. By conjugating a lauric acid chain to a repetitive IIRR motif, CIR self-assembles into protease-resistant nanofibrils, forming a hydrogel that enables sustained intestinal release following oral administration. The hydrogel exhibits a structural transformation triggered by liposaccharide (LPS), a pro-inflammatory molecule produced by pathogenic bacteria at UC loci. Physiologically stable β-sheet can transfer to bioactive α-helix conformations, promoting localized therapeutic efficacy and circumventing systemic toxicity. In murine models of acute and chronic colitis, orally administered CIR hydrogel demonstrated better performance than 5-ASA. The LPS binding peptide selectively destroy pathogenic bacteria, simultaneously silencing TLR4-driven cytokine storms. Through this coordinated action, CIR resolves the intertwined immune-microbial pathology, offering a safe and effective strategy for UC management. Our work highlights the potential of self-assembly peptide hydrogels as next-generation therapeutics for complex inflammatory disorders.

## Results

### Rationally Engineered HDP Self-Assembles into Nanofiber Hydrogels

To address the limitations of natural HDPs, we designed C12-(IIRR)₂I-NH₂ (CIR), an arginine-rich peptide featuring an alternating hydrophobic/cationic (XXYY) motif to drive α-helix formation and microbial interactions (Fig. 1a and Extended Data Fig. 1a–c).^22–24^ While α-helical structures often confer antimicrobial activity, their inherent cytotoxicity and instability prompted us to modify the peptide with a lauric alkyl chain. This substitution induced a structural shift to β-sheets, which are more stable under physiological conditions and facilitate self-assembly into nanofibers. Critically, the β-sheet conformation remains inert until encountering pathogenic bacteria, where LPS-binding triggers a structural transformation to the bioactive α-helix. Arginine residues were strategically incorporated to enhance peptidase resistance and immunomodulatory capacity.^25–27^

**Fig. 1.**
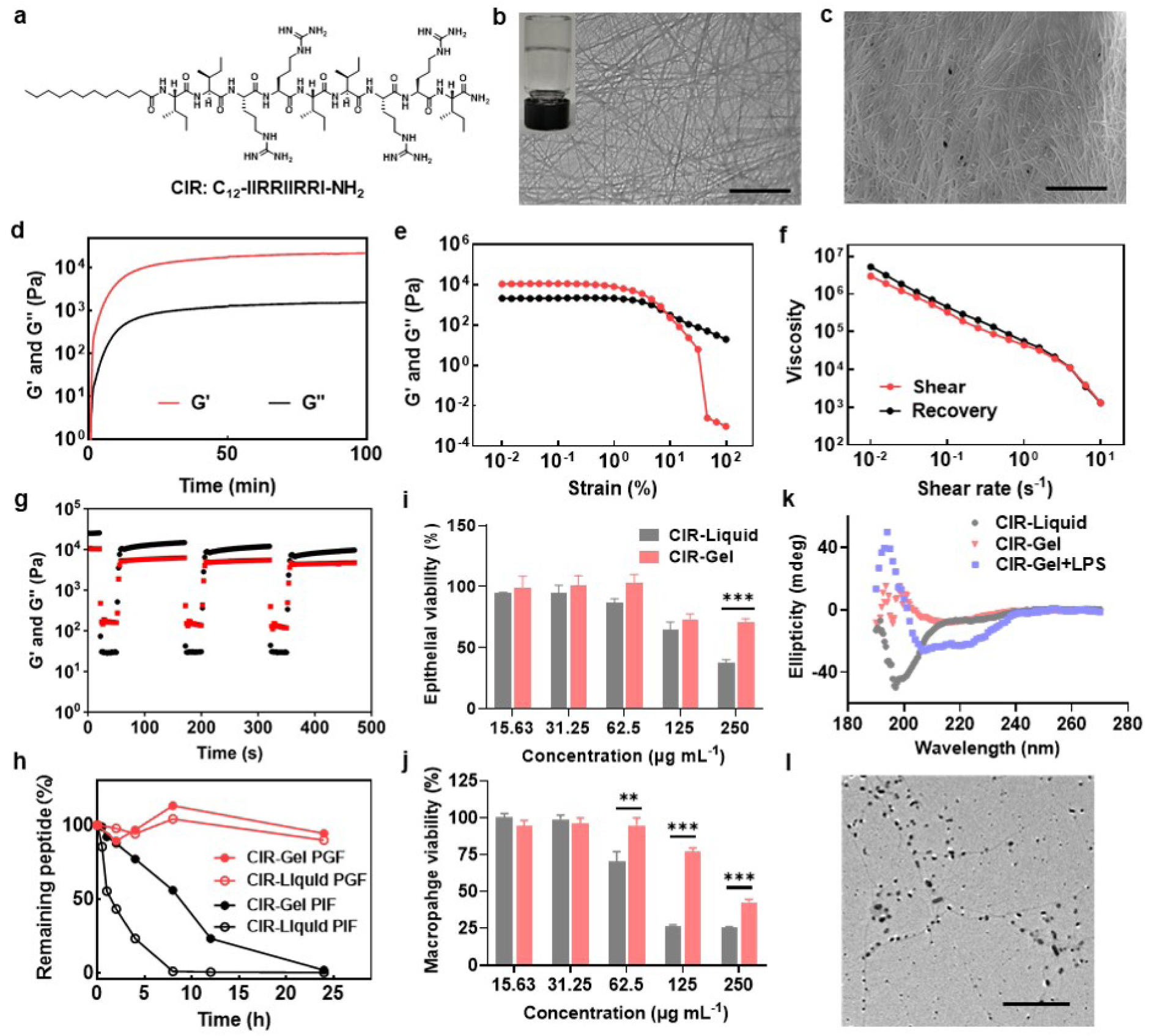
Rationally engineered HDP self-assembles into nanofiber hydrogels with oral-compatible stability and structural switching property. **a,** Chemical structure of the rationally designed HDP with an alternating hydrophobic/cationic motif and a lauric alkyl substitution. **b,c**, The nanofibrillar network of the CIR hydrogels as captured by TEM (b) and cryo-SEM (c). Scale bar, 2 μm. Insert: a photograph of inverted CIR hydrogels. **d**, Rapid gelation of CIR hydrogels as measured by a rheometer. G’, storage modulus; G’’, loss modulus. Final CIR concentration was 5 mg·mL^-1^, with addition of 10 mM phosphate buffer and 50 mM Tris at pH 7. **e-g**, Rheological analysis of CIR hydrogels with strain sweep (e), continuous flow (f) and cyclic strain time sweep (g), demonstrating that CIR self-assembled into hydrogels with robust mechanical strength, shear-thinning and self-healing property. **h**, Degradability of the CIR hydrogel and liquid in UC patient gastric fluid (PGF) and intestinal fluid (PIF), showing improved stability of CIR after gelation. **i,j**, Cell viability assessment of CIR liquid and hydrogels in NCM460 (i) and RAW264.7 (j) as measured by MTT, which showed reduced cytotoxicity after forming hydrogel. **k**, CD spectrum showing the secondary structure transformation of CIR from the stable β-sheet into bioactive α-helix with the addition of LPS. **l**, TEM figure of CIR hydrogels after the addition of LPS exhibiting the degradation of nanofiber structure. Scale bar, 2 μm. Data represent means ± SEM. * p < 0.05, ** p < 0.01, *** p < 0.001, by Student’s t test.

The CIR peptide exhibited rapid, buffer-dependent hydrogelation, forming a stable matrix at concentrations as low as 0.1 mg ml^-1^. Transmission and cryo-scanning electron microscopy (TEM/cryo-SEM) confirmed a three-dimensional nanofibrillar network (Fig. 1b,c). Gelation occurred within 30 min, driven by hydrogen bonding and hydrophobic interactions (Fig. 1d). Time-resolved fluorescence assays using Thioflavin T (ThT) and Nile red revealed sequential stabilization: hydrogen bonds dominated initial gelation (equilibrium at 100 min), followed by hydrophobic rearrangement during aging (equilibrium at 200 min) (Extended Data Fig. 1d).^28,29^ These dynamic interactions underpinned the hydrogel’s mechanical robustness, enabling its use in physiologically relevant environments.

### CIR Hydrogel Demonstrates Oral Delivery-Compatible Rheological and Stability Profiles

The rheological properties of CIR hydrogel were characterized by oscillatory shear measurements. Storage (G′) and loss (G′′) moduli revealed concentration-dependent mechanical strength, plateauing at 5–10 mg ml^-1^ (Fig. 1e and Extended Data Fig. 1e). The hydrogel displayed shear-thinning behavior with viscosity decreasing under high shear rates, and full recovery of G′/G′′ after cyclic strain, confirming robust self-healing capacity (Fig. 1f,g). These traits enable easy syringeability for oral dosing and rapid reformation in the gastrointestinal tract.

A key hurdle for peptide therapeutics is gastric degradation. However, the molecular design of CIR circumvents this problem. The cationic charge, low molecular weight, and absence of pepsin cleavage sites due to arginine termini together contribute to its high stability in acidic environment. In simulated gastric fluid (SGF, pH 1.2), both liquid and hydrogel forms remained intact for more than 24 hours (Extended Data Fig. 1f). While intestinal proteases degraded free CIR in simulated intestinal fluid (SIF), hydrogelation slowed degradation and extended half-life. This stability was recapitulated in gastric and intestinal fluids from UC patients, confirming translational relevance (Fig. 1h).

### CIR Hydrogel Enhances GI Retention and Mitigates Cytotoxicity via Structural Transformation

To evaluate gastrointestinal (GI) retention, we orally administered indocyanine green (ICG)-labeled CIR hydrogel to dextran sulfate sodium (DSS)-treated mice with UC. In vivo fluorescence imaging revealed prolonged GI retention, with signal enrichment in the abdominal region beginning at 4 hours post-administration. Ex vivo analysis at 24 hours confirmed the enhanced retention and accumulation in colon tissues, which resulted from the hydrogelation (Extended Data Fig. 1g-j).

The therapeutic potential of CIR hydrogel was further underscored by its reduced cytotoxicity. In vitro assays using colon epithelial (NCM460) and macrophage (RAW 264.7) cell lines showed minimal toxicity, owing to its β-sheet-dominated structure under physiological conditions (Fig. 1i,j). This inert conformation masked the bioactivity of CIR peptide until encountering LPS, a key component from pathogenic bacteria that can induce excessive inflammation in colon tissues. The peptide hydrogel consequently disassembled and restored the antimicrobial α-helix conformation (Fig. 1k,l). This LPS-triggered transformation ensures targeted activity at infection sites while sparing probiotics and host cells.

### CIR Hydrogel Alleviates Colitis Symptoms in Both Acute and Chronic Colitis Mouse Model

In a DSS-induced acute colitis murine model, oral administration of CIR hydrogel robustly mitigated disease severity (Fig. 2a). CIR-treated mice exhibited minimal weight loss, preserved colon length, and near-complete resolution of diarrhea and hematochezia (Fig. 2b-d), better than the clinical standard 5-ASA in colon restoration (Fig. 2e). Histological analysis confirmed the protective effects of CIR. Colonic architecture in the CIR group was similar to healthy controls, with intact crypts, minimal ulceration, and reduced immune infiltration, whereas 5-ASA showed partial recovery (Fig. 2f). The hydrogel also restored mucosal barrier integrity, elevating tight junction proteins Claudin-1 and ZO-1 to near-normal levels, which is a critical factor in mitigating systemic inflammation (Fig. 2g-i).

**Fig. 2.**
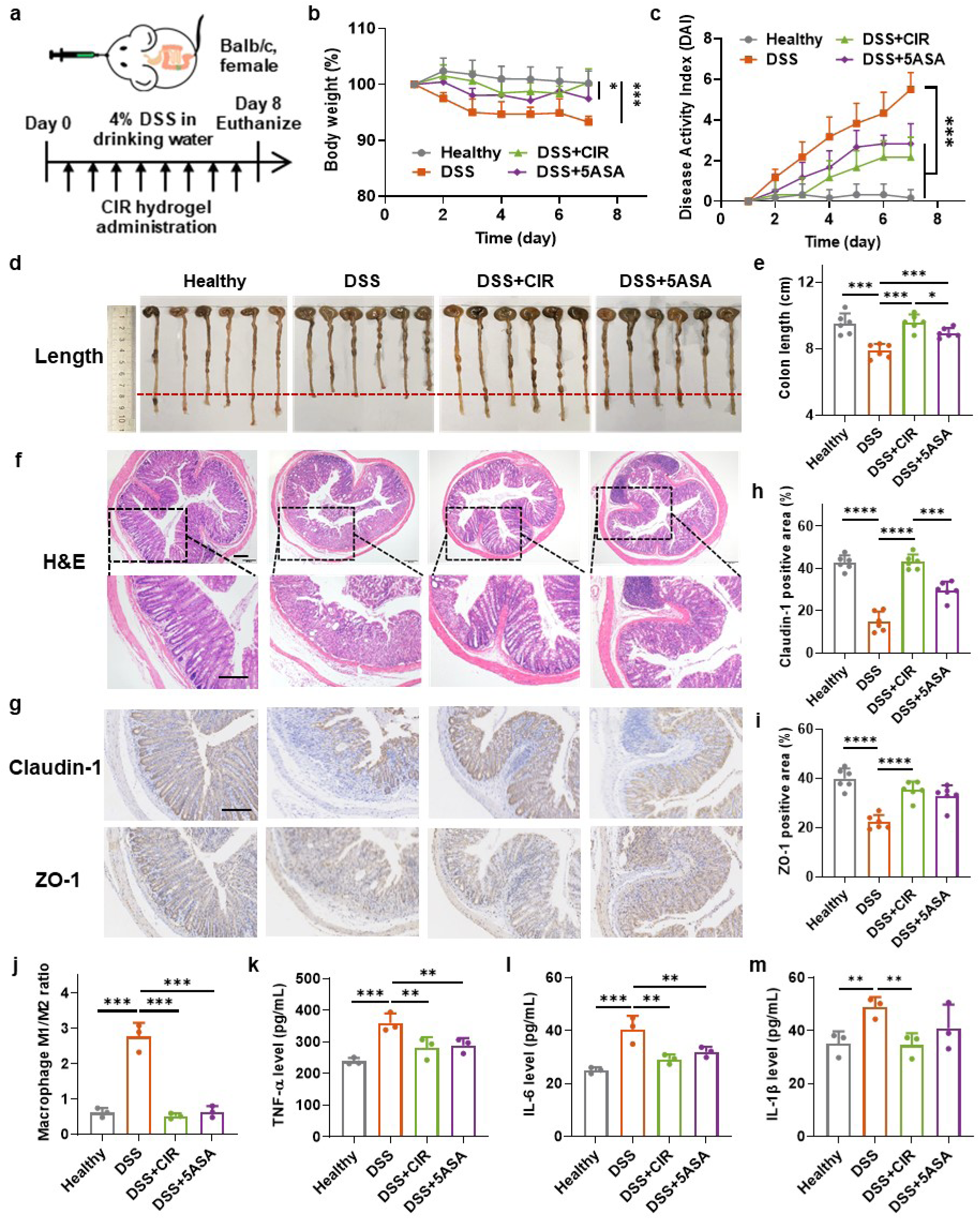
CIR hydrogels alleviated colitis symptoms and reduced inflammation in acute colitis mouse model. **a,** Experimental design of CIR hydrogel treatment in a DSS-induced acute colitis mouse model. **b,c,** Daily changes of body weights (**b**) and DAI scores (**c**) of mice showing alleviated colitis symptom under CIR hydrogel treatment. **d,e,** Colon photos (**d**) and lengths (**e**) of mice after sacrificed on day 8 exhibiting increased colon length in CIR hydrogel group. **f,** H&E staining images for pathological diagnosis of colon tissues demonstrating improved colon tissue morphology under CIR hydrogel treatment. Scale bar, 200 μm. **g,** IHC staining images for Claudin and ZO-1 of colon tissues showing higher expression levels of tight junction proteins in CIR hydrogel group. Scale bar, 200 μm. **h,i,** Quantification of positive staining area of Claudin (h) and ZO-1 (i) in colon tissue sections. **j,** M1/M2 ratio of macrophages extracted from mouse spleens as measured by flow cytometry demonstrated that CIR hydrogel induced macrophage differentiation to M2 subtype. **k-m,** The levels of TNF-α (k), IL-6 (l), and IL-1β (m) in mouse serum as determined by ELISA assay indicated that CIR hydrogel inhibited the release of inflammatory cytokines. Data represent means ± SEM. * p < 0.05, ** p < 0.01, *** p < 0.001, **** p < 0.0001, by Student’s t test.

The immunomodulatory potency of CIR was evidenced by macrophage polarization both in vitro and in vivo. In LPS-stimulated mouse bone marrow-derived macrophages (BMDMs) and RAW 264.7 cell line, CIR treatment reduced the M1/M2 ratio, as quantified by flow cytometry analysis of surface markers CD80 and CD206, representing subtype M1 and M2, respectively (Extended Data Fig. 2a-d). In murine splenic tissues, flow cytometry also revealed a dose-dependent reduction in the pro-inflammatory M1/M2 ratio (Fig. 2j and Extended Data Fig. 2e,f), paralleled by decreased serum levels of TNF-α, IL-6, and IL-1β (Fig. 2k-m). This anti-inflammatory activity resulted from a combinatorial effect of LPS binding and macrophage metabolism reprogramming, which will be explained in detail later.

Safety profiling confirmed the biocompatibility of CIR hydrogel. No histopathological abnormalities in major organs or perturbations in hepatic/renal serum markers (ALT, AST, LDH, CREA, and urea) were observed (Extended Data Fig. 3a,b), underscoring its translational viability.

To address the relapsing nature of UC, we evaluated the therapeutic effect of CIR hydrogel in a chronic colitis murine model mimicking cyclical inflammation (Fig. 3a). Mice subjected to three DSS cycles exhibited severe colon shortening, crypt damage, and barrier disruption, which were all markedly reversed by CIR treatment (Fig. 3b-e). Notably, CIR surpassed 5-ASA in restoring colon length and tight junction protein expression. The performance was even better than that in the acute colitis model, underscoring its superiority in sustained disease management.

**Fig. 3.**
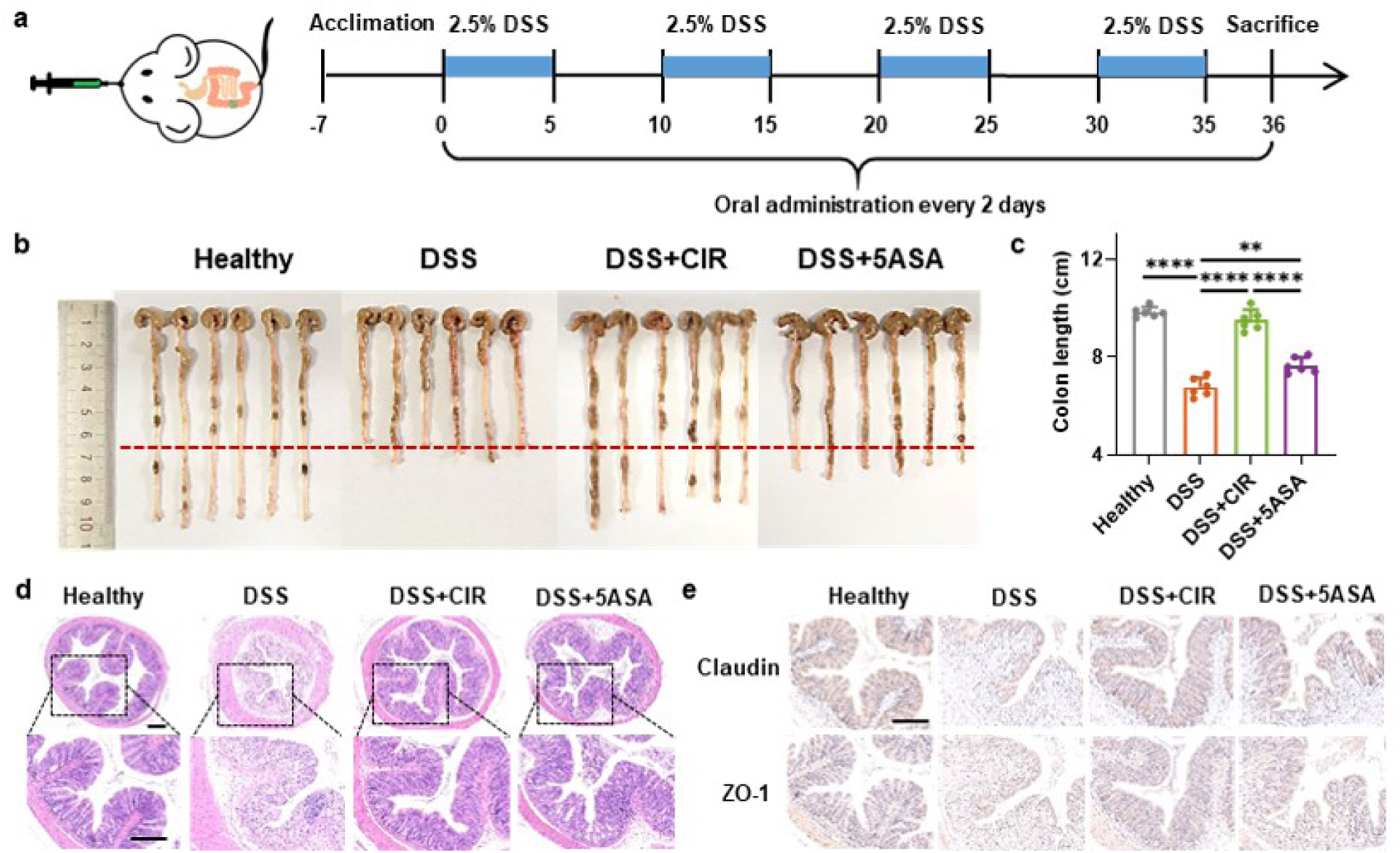
CIR hydrogels alleviated colitis symptoms in chronic colitis mouse model. **a,** Experimental design of CIR hydrogel treatment in a DSS-induced chronic colitis mouse model. **b,c,** Colon lengths (b) and representative photos (c) of mice receiving various treatment after sacrificed on day 36. **d,** H&E staining images for pathological diagnosis of colon tissues. Scale bar, 200 μm. **e,** IHC staining images for Claudin and ZO-1 of colon tissues. Scale bar, 200 μm. Data represent means ± SEM. * p < 0.05, ** p < 0.01, *** p < 0.001, **** p < 0.0001, by Student’s t test.

### CIR Hydrogel Suppresses Colitis via LPS-Activated Restoration Effect of Microbiota Homeostasis

Given its promising therapeutic efficacy in colitis models, elucidating the mechanism of CIR hydrogel is essential. Gut microbiota dysbiosis, marked by diminished diversity and pathogenic enrichment, drives UC progression. This effect mainly results from pathogenic bacteria continuously producing LPS that induces inflammation in colon. A LPS-triggered HDP can be a promising candidate to treat these pro-inflammatory pathogens. To evaluate the microbiota modulation ability of CIR hydrogel, we performed 16S rRNA sequencing on fecal samples from murine and NHP colitis models. Operational taxonomic unit (OTU) analysis revealed that CIR-treated groups shared greater microbial overlap with healthy controls than DSS groups (Fig. 4a), indicating a microbiota rebalance under treatment. In both mouse models, CIR restored α-diversity indices (Shannon, Simpson, Pielou_e) and β-diversity to near-healthy levels (Fig. 4b,c), further demonstrating the microbiota rebalancing effect, whereas 5-ASA showed little beneficial effect and even exacerbated dysbiosis in β-diversity analyses (Extended Data Fig. 4a,b).

**Fig. 4.**
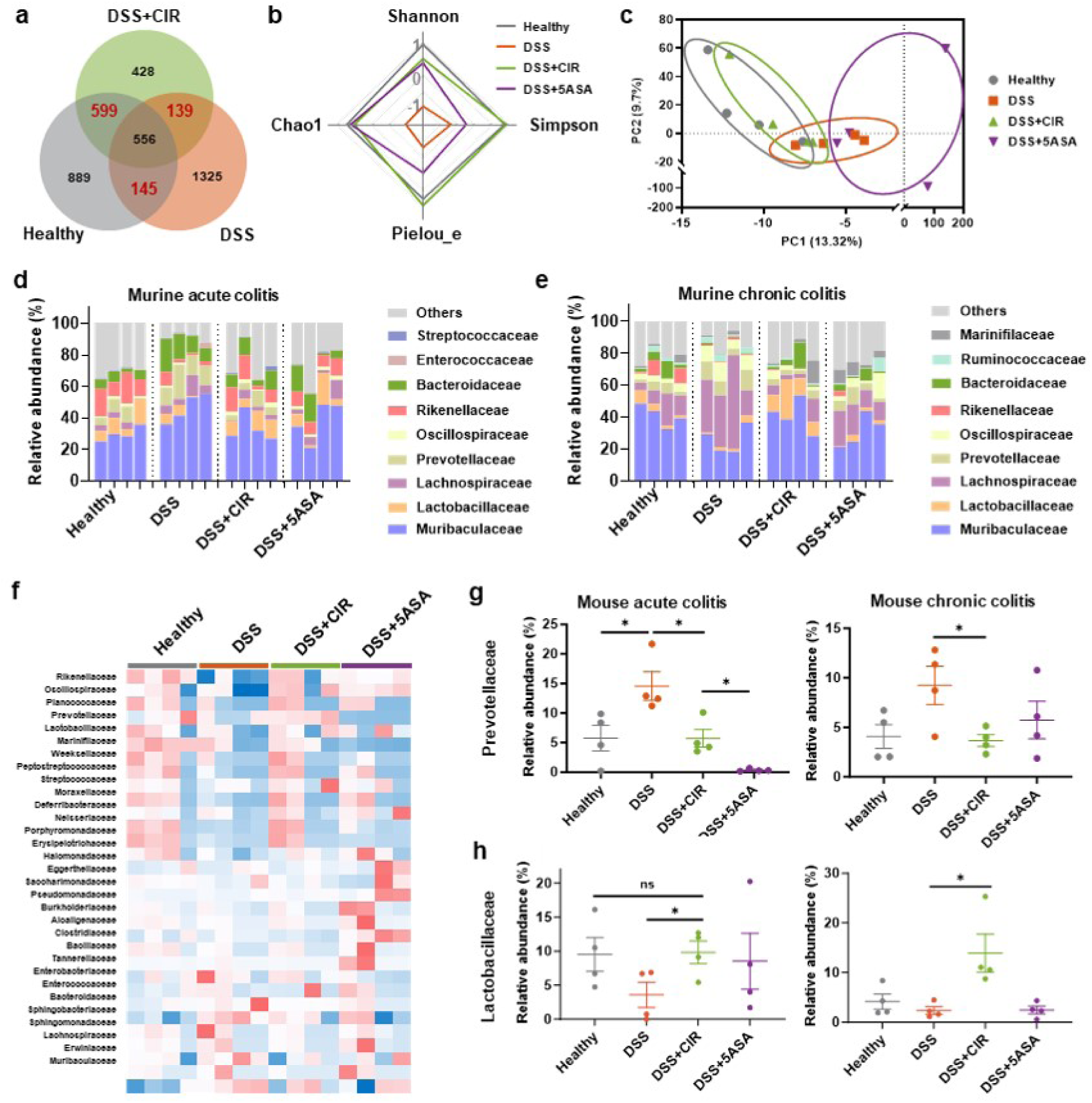
CIR hydrogels modulated microbiota abundance and diversity in colitis mouse models. **a,** Venn diagram showing the number of unique and overlapped OTUs among Healthy, DSS, and DSS + CIR groups. **b,** Radar diagram showing α diversity with Shannon, Simpson, Pielou_e, and Chao1 index. **c,** PCA figure showing the *β*-diversity, each point represents one mouse (n = 4). **d,e**, Relative abundance of microbiota that is significantly altered at family levels in acute colitis (d), and chronic colitis (e) mouse models. **f,** Heatmap of microbiome in mouse model displaying the different microbial abundance in family level among groups. **g,h,** The abundance changes of *Prevotellaceae* (g) and *Lactobacillaceae* (h) among different groups showing similar trends in all three models. Data represent means ± SEM. * p < 0.05, ** p < 0.01, *** p < 0.001, by Student’s t test.

At the phylum level, CIR normalized the Bacteroidota/Firmicutes ratio in both models by suppressing LPS-producing Bacteroidota and enriching SCFA-generating Firmicutes (Extended Data Fig. 4c,d). Family-level profiling highlighted CIR’s dual action: depleting pathobionts like *Prevotellaceae* that is one of the main source of LPS in UC patients, and enriching probiotics such as *Lactobacillaceae* that can modulate macrophage activity via S100a8/TLR pathways (Fig. 4d,e). Heatmap clustering confirmed that CIR-treated microbiota mirrored healthy states (Fig. 4f and Extended Data Fig. 4f). Under CIR hydrogel treatment, reduced *Prevotellaceae* amounts were observed in all models, consistent with the hypothesized LPS-triggered antimicrobial effect (Fig. 4g). Some other LPS producing pathogens were also inhibited, such as *Bacteroidaceae*, *Lachnospiraceae* and *Spirochaetaceae*. In contrast, probiotics like *Lactobacillaceae* lack LPS on their membranes, which were less affected by CIR treatment and exhibited increased proportions in these models (Fig. 4h). *Lactobacillaceae* is known for its lactic acid production and anti-inflammatory effects, which would promote the recurrence of UC.

### CIR Hydrogel Mitigates Abnormal Inflammation by Downregulating TLR4 Related Pathways

In addition, LPS level change from microbiota rebalance would contribute to the regulation of inflammation and immune environment. RNA sequencing (RNA-seq) of mouse colon tissues revealed that inflammatory pathways such as TLR, MAPK, and JAK-STAT were upregulated in the DSS group but downregulated in the DSS+CIR group, demonstrating the capacity of CIR peptide to remodel the immune microenvironment at UC sites (Extended Data Fig. 5a-d). Heatmap revealed that representative genes in these pathways, such as S100a9, Fcgr1 and Tlr4, were downregulated, combined with protein-protein interaction (PPI) network analysis identifying TLR4 as a key player in inflammatory bowel disease (IBD) pathway (Fig. 5a and Extended Data Fig. 5e). Elevated expression of Tlr4 and Il18r1 in the DSS group was significantly reduced by CIR hydrogel (Data Fig. 5b,c).

**Fig. 5.**
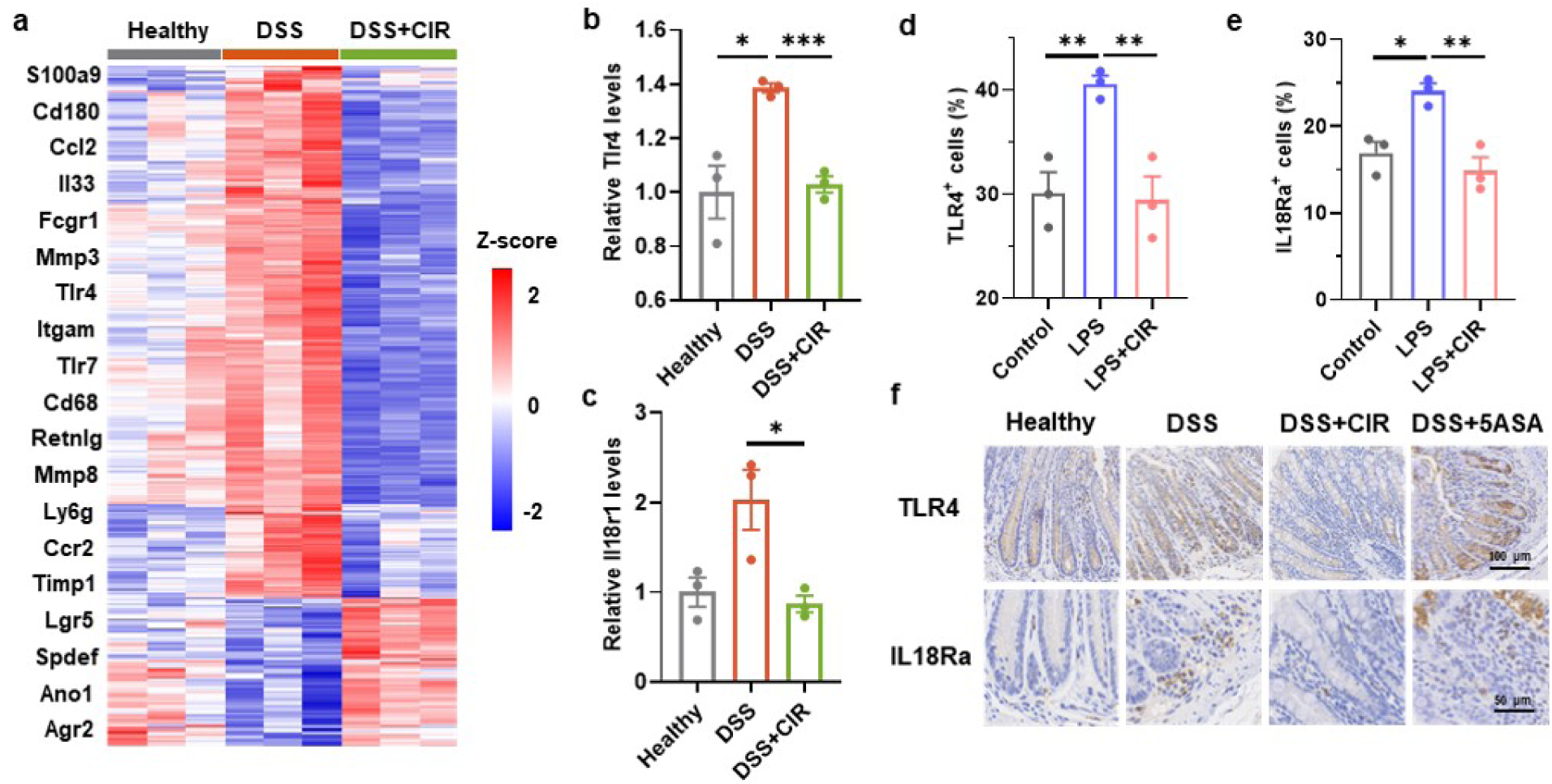
CIR hydrogels deactivated abnormal inflammation and immune response by suppressing TLR4-related pathway. **a,** Heatmap of differentially expressed genes in the UC mouse model showing a similar transcriptomic profile between DSS+CIR and Healthy groups. **b,c,** Transcript levels of Tlr4 (b) and Il18r1 (c) downregulated by CIR hydrogel treatment. **d,e,** Quantification of TLR4^+^ (d) and IL18Ra^+^ (e) cells in BMDM from flow cytometry results. **f,** IHC staining images for TLR4 and IL-18Ra of colon tissues under different treatments. Data represent means ± SEM. * p < 0.05, ** p < 0.01, *** p < 0.001, by Student’s t test.

Functional validation using LPS-induced M1-polarized bone marrow-derived macrophages (BMDMs) showed decreased proportions of TLR4⁺ and IL-18Rα⁺ cells and consistent RNA level changes of related genes under CIR treatment (Fig. 5d,e and Extended Data Fig. 5f,g). Immunohistochemistry (IHC) of colon sections from colitis mice further confirmed the downregulation of TLR4 and IL-18 receptor levels, outperforming 5-ASA (Fig. 5f).

## Discussion

We present an orally administered self-assembled peptide hydrogel, CIR, that synergistically targets immune dysregulation and gut microbiota dysbiosis to achieve durable remission in UC. By leveraging rational design principles inspired by HDPs, CIR combines structural adaptability, biocompatibility, and dual bioactivity to address critical gaps in current UC therapies. Its β-sheet nanofiber architecture ensures stability during gastrointestinal transit, while microenvironment-responsive switching to α-helix conformations at inflamed sites enables localized therapeutic action. This design minimizes enzymatic degradation and systemic toxicity, persistent challenges for peptide-based drugs, while maximizing bioavailability at disease loci.

Conventional murine acute colitis models remain essential for therapeutic screening, yet their translational limitations stem not only from interspecies physiological divergences, but also from inadequate modeling of UC’s chronic nature.^32^ In this study, we utilized chronic colitis mouse and NHP models with multi-cycle DSS exposure, aligning with human UC’s relapsing-remitting course, which bridged translational gaps through gastric secretion parallels and microbiota homology. This cross-species validation strategy demonstrated the therapeutic superiority of CIR hydrogel, underscoring its translational potential as a clinically viable UC therapeutic.

In murine models of acute and chronic colitis, CIR outperformed the clinical standard 5-ASA by restoring mucosal integrity, suppressing pro-inflammatory cytokines, and normalizing colon function. Mechanistically, 16S rRNA sequencing demonstrated CIR’s capacity to rebalance gut microbiota, enriching probiotic Lactobacillaceae, known for lactic acid production and macrophage modulation; while depleting pathobiont Prevotellaceae, a main source of LPS. Concurrently, transcriptomic studies revealed that CIR silenced TLR4-driven hyperinflammation by LPS binding, thereby attenuating downstream JAK-STAT/IL-18 signaling, which regulated neutrophil recruitment and macrophage metabolism. The LPS-triggered structure switching enabled the CIR peptide to selectively destroy those pro-inflammatory pathogenic bacteria, and the rebalanced microbiota facilitates the restoration of immune homeostasis. By rectifying dysbiosis while silencing immune hyperactivation, CIR disrupts the bacteria-immune feedback loop central to UC. Its superior efficacy over 5-ASA, which lacks microbiota-modulating activity, positions it as a dual-action therapeutic addressing both inflammation and microbial imbalance.

The structural and functional advantages of CIR address two unmet needs in UC therapy. Firstly, clinical therapeutics (e.g., 5-ASA, biologics) lack microbiota-modulating activity, failing to resolve intertwined immune-microbial pathology. Secondly, systemic immunosuppressants suffer from drug resistance and off-target risks, not suitable for long-term treatment. By contrast, the CIR self-assembled hydrogel circumvent these limitations through localized synergistic bioactivity and oral delivery stability, offering a paradigm shift toward precision therapeutics.

While this study establishes preclinical proof-of-concept, future work should explore long-term safety, dose optimization, and human microbiota interactions. Nevertheless, CIR’s ability to concurrently resolve immune and microbial dysfunction positions it as a prototype for multifunctional therapies in inflammatory diseases. By bridging peptide engineering, immunology, and microbiome science, this work opens avenues for next-generation biologics tailored to complex, multifactorial disorders.

## Methods

### Materials

The CIR peptide (>98% purity) was purchased from Shenzhen JYMed Technology Co., Ltd (China). Buffer and salts were sourced from Energy Chemicals Co., Ltd (China) and cell culture media from BioLegend Company (China). Lipopolysaccharide (LPS) was obtained from Sigma-Aldrich (USA), while IL-6 and TNF-α ELISA kits were purchased from ExCell Bio (China). RAW 264.7 murine macrophage and NCM460 human colon mucosal epithelial cell lines were obtained from the Chinese Academy of Medical Sciences. Deionized water was generated from a Milli-Q Plus water purification system.

### Preparation of the CIR hydrogel

CIR peptide powder was dissolved in deionized water at concentrations of 2, 4, 10, and 20 mg ml^-1^, followed by 30 s ultrasonication and incubation at 80 °C until complete dissolution. A buffer solution (pH 7.4) containing 20 mM phosphate and 100 mM Tris was prepared, which was then mixed with the peptide solution at a 1:1 (v/v) ratio. The mixture was aged for 72 h at 25 °C to form hydrogel. A final hydrogel concentration of 5 mg ml^-1^ was used for further study.

### Morphological analysis

For TEM imaging. The peptide hydrogel was diluted 10-fold with buffer, and deposited onto 300-mesh copper grids. Images were acquired using a JEM-1011 TEM system (JEOL, Japan). Cryo-SEM samples were prepared by flash-freezing hydrogel-coated silicon substrates in liquid nitrogen, followed by sublimation in a freeze-dryer and sputter-coating with 5 nm platinum. Images were obtained on a S-4300 SEM system (Hitachi, Japan).

### Spectroscopic characterization

Fluorescence spectra were recorded on an FLS1000 spectrometer (Edinburgh Instruments, UK). Thioflavin (ThT, 10 μM) or Nile red (10 μM) was incorporated into 5 mg ml^-1^ CIR hydrogels, with excitation wavelengths set to 440 nm and 477 nm, respectively. Circular dichroism (CD) spectra were measured on a J-1700 spectrometer (JASCO, Japan). The peptide aqueous solutions (5 mg ml^-1^) were diluted 100-fold with ultrapure water or lipopolysaccharide (LPS) solution, while the peptide hydrogels (5 mg ml^-1^) were diluted 100-fold using PBS or LPS-supplemented PBS. All samples were transferred into 1 cm quartz cuvettes and subjected to CD with a scan range of 190– 270 nm.

### Rheological analysis

An MCR302 rheometer (Anton Paar, Austria) was employed for rheological analysis. Dynamic time sweeps monitored storage modulus (G’) and loss modulus (G’’) evolution over 100 min. Frequency sweeps (0.1-10 rad·s^-1^) and strain sweeps (0.01-100%) were performed at 20 °C. Sample viscosity was cyclically measured within the shear rate range of 0.01-10 s^-1^. Recovery behavior was assessed under repeated strain switch between 1% and 100% s. Rheological experiments.

### Stability evaluation

The peptide aqueous solutions and hydrogels (5 mg ml^-1^) were diluted 100-fold with artificial gastric fluid (prepared with 16.4 mL diluted hydrochloric acid and 10 g pepsin per liter, pH 1.2–1.5) or artificial intestinal fluid (prepared with 6.8 g KH_2_PO_4_ and 10 g pancreatin per liter, pH 6.8), then incubated at 37°C. Aliquots were collected at 0, 0.5, 1, 2, 4, 8, 12, and 24 hours, mixed with acetonitrile and water (1:1 v/v), filtered through 0.22 μm membranes, and analyzed via an UltiMate 3000 HPLC system (Thermo Fisher, USA) using a C18 column. The mobile phase consisted of 0.1% trifluoroacetic acid in water (A) and 0.1% trifluoroacetic acid in acetonitrile (B), with a gradient program: 50% B (0–16 min), 85% B (16–19 min), and re-equilibration to 50% B. UV detection was performed at 214 nm with a flow rate of 0.6 mL/min. Patient-derived gastric/intestinal fluids were processed identically.

### Cell viability assay

NCM460 and RAW 264.7 cells were cultured in DMEM medium containing 10% fetal bovine serum and 1% penicillin-streptomycin antibiotics at 37 °C under a 5% CO_2_ atmosphere. Cytotoxicity of peptides was evaluated using a standard MTT assay. Cells were seeded in 96-well plates (10^4^ cells/well) and allowed to adhere for 24 h. After removing the culture medium, peptide solutions were added and incubated for 24 h. Subsequently, 0.5 mg ml^-1^ MTT reagent was added into each well and incubated for 4 h to form formazan crystal, followed by dissolution with 100 μl DMSO and measurement of absorbance at 570 nm using a microplate reader.

### DSS induced colitis mouse model

Female Balb/c mice (6–8 weeks old, 18 ± 2 g) were purchased from the Vital Laboratory Animal Center (Beijing, China). Mice were acclimatized under standard laboratory conditions (22°C, 12 h light/dark cycle) for 7 days prior to experiments. Acute colitis was induced by administering 4% DSS in drinking water for 7 consecutive days. Mice were randomized into four groups (n = 6): (1) Healthy, (2) DSS + 100 μl PBS (DSS), (3) DSS + 25 mg kg^-1^ CIR hydrogel (DSS+CIR), (4) DSS + 25 mg kg^-1^ 5-ASA (DSS+5ASA). Therapeutic agents or PBS were administered daily via oral gavage. Disease activity index (DAI) scores were calculated based on weight loss, stool consistency, and fecal blood. On day 8, mice were euthanized, followed by collections of major organs (heart, liver, spleen, lung, and kidney). Colons were harvested for histopathology and transcriptomic profiling. Fecal samples collected on day 7 were analyzed for microbial composition via 16S rRNA sequencing. For chronic colitis model, mice underwent three cycles of 2.5% DSS administration (5 days DSS + 5 days recovery per cycle), followed by a final five days of 2.5% DSS solution. Therapeutic agents or PBS were administered every other day. Colon tissues and fecal samples were processed identically to the acute model. All animal experiments were approved by the Experimental Animal Ethics Committee of the Institute of Process Engineering, Chinese Academy of Sciences (Approval No. IPEAECA2023051).

### Biodistribution study

Female Balb/c mice were randomized into two experimental groups (n = 3): DSS + free ICG and DSS + CIR-ICG. Colitis was induced via ad libitum administration of 4% DSS in drinking water. On day 7, mice received 100 μL hydrogel (5 mg ml⁻¹ CIR + 0.2 mg mL⁻¹ ICG) or free ICG solution via oral gavage. In vivo fluorescence imaging (Ex/Em: 745/820 nm) was performed at 0, 1, 2, 4, 8, 12, and 24 h post-administration using IVIS imaging system (PerkinElmer, USA). At 24 h, mice were euthanized, and major organs (heart, liver, spleen, lung, kidney, stomach, small intestine, and colorectal tissue) were excised for ex vivo imaging. Colorectal specimens were washed with ice-cold PBS to remove fecal contents prior to secondary imaging. Fluorescence intensity was quantified using Living Image software with identical exposure settings across samples.

### Immunohistochemistry

Paraffin-embedded tissue sections underwent antigen retrieval in 0.01 M citrate buffer (pH 6.0) at 95°C for 20 minutes. After blocking with 5% serum, endogenous peroxidase was quenched using 3% H₂O₂ solution. Overnight primary antibody incubation (4 °C) was performed using validated monoclonal antibodies Post-incubation, sections were washed and labeled with species-matched secondary antibodies, and signal amplification was achieved through chromogenic detection using Pierce DAB Substrate Kit (Thermo Fisher, USA). Counterstaining with hematoxylin was performed to delineate cellular nuclei. Stained slides were digitized using the Aperio Digital Pathology Slide Scanner (Leica, USA), yielding high-resolution brightfield images. Quantitative analysis of immunostaining intensity was conducted using ImageJ software (National Institutes of Health, Bethesda, MD, USA) with color deconvolution algorithms to separate chromagenic signals from background artifacts. Image acquisition parameters and analytical settings were standardized across all experimental replicates to ensure reproducibility. Tissue sections were stained with the following antibodies according to the manufacturer’s instructions: from Servicebio: Claudin (GB15032), ZO-1 (GB115686), TLR4 (GB15186), IL18R (GB114098).

### Flow cytometry

BMDM cells were isolated from female Balb/c mice (6-8 weeks old). Femurs and tibias were excised and immersed in 75% ethanol for 5 minutes. Bone marrow cavities were flushed thoroughly with PBS, and marrow cells were collected through a 70-μm cell strainer. The cell suspension was centrifuged at 1000 g for 5 min to remove debris. Red blood cell lysis solution was added and incubated on ice for 10 minutes, followed by centrifugation at 1000 g for 5 min to obtain purified BMDM cells. Cells were cultured in RPMI1640 supplemented with 20 ng/mL macrophage colony-stimulating factor (M-CSF) and 10% FBS at 37°C in 5% CO₂. Medium replacement occurred every 48 hours until day 6, when cells reached mature macrophage differentiation. On day 7, cytokine-free RPMI1640 was employed for subsequent experiments. To induce M1 polarization, BMDM cells were treated with 100 ng/mL LPS and 5 ng/mL interferon-gamma (IFN-γ) for 24 hours. Post-polarization induction, cells were co-cultured with CIR hydrogel in different concentrations for an additional 24 hours. Cellular polarization status was assessed via flow cytometry (BD FACSAria™ Fusion, BD Biosciences, USA).

Spleens were harvested from euthanized mice in colitis model and minced gently on ice using a sterile razor blade in RPMI-1640 medium containing 2% FBS. The tissue homogenate was passed through a 70-μm cell strainer to obtain single-cell suspensions. Cells were centrifuged at 1000 g for 5 min, and the supernatant was discarded. Red blood cell lysis was performed by resuspending the pellet in 1 mL lysis buffer for 3 min on ice, followed by neutralization with 2 mL RPMI-1640 medium. The resultant cells were washed twice and resuspended in RPMI-1640 medium containing 1% FBS for subsequent analysis. Cell viability was determined using Trypan Blue exclusion (Invitrogen, USA), and viable cells were harvested for flow cytometric polarization assessment. Cells were stained with the following antibodies according to the manufacturer’s instructions: from invitrogen, F4/80 (FITC), CD80 (PE), CD206 (PerCP-eFluor710), TLR4 (Alexa FluorTM488), IFN-γ (APC), IL18Rα (PE).

### ELISA

Viable cells from mouse spleens were seeded at 2 × 10⁵ cells/well in 6-well plates. After 24-hour incubation, supernatants were collected for cytokine analysis. TNF-α, IL-6, and IL-1β levels in supernatants were quantified using ELISA kits following manufacturer protocols: capture antibody immobilization, blocking with 5% BSA, sample incubation, and detection using HRP-conjugated antibodies with TMB substrate, and terminated by 2 M H₂SO₄. Optical density at 450 nm was measured using a microplate reader, and concentrations were calculated based on standard curves. RAW 264.7 macrophages (10^4^ cells/well) were cultured in 48-well plates for 24 h. Cells were stimulated with 1 μg ml^-1^ LPS in the absence or presence of peptides. Supernatants collected at 24 h post-stimulation were centrifuged at 300 g for 5 min to remove debris.

### RNA sequencing and analysis of mouse colon tissues

Total RNA was also extracted from mouse colon tissues and evaluated for integrity using the Bioanalyzer 2100 system. Polyadenylated mRNA was enriched via polyA selection with magnetic beads and then chemically fragmented. First-strand cDNA synthesis was carried out with random hexamer primers and M-MuLV reverse transcriptase. After RNA removal, second-strand cDNA was synthesized using DNA polymerase I. The cDNA products underwent end repair, polyA tailing, and adapter ligation. Fragments of 370–420 bp were size-selected with AMPure XP beads, amplified by PCR, and purified to generate final libraries. Cluster generation was performed on an Illumina cBot system with the TruSeq PE Cluster Kit v3, followed by 150 bp paired-end sequencing on the NovaSeq 6000 platform.

Differential gene expression across experimental conditions was assessed using DESeq2 (version 1.20.0), with statistical significance defined as Benjamini–Hochberg adjusted p-values ≤ 0.05. Functional annotation of DEGs was performed via GO and KEGG pathway enrichment analyses using the clusterProfiler package, which included correction for gene length bias. Protein–protein interaction networks were constructed using the STRING database (version 11.5), incorporating both experimentally verified and in silico predicted interactions.

### 16S rRNA sequencing of gut microbiota

Faecal samples were collected and DNA was extracted from single faecal pellets using the Promega Maxwell RSC PureFood GMO and Authentication Kit (Promega, USA) according to the manufacturer’s instructions. The V3-V4 hypervariable regions of bacterial 16S rRNA genes were amplified using barcoded primers (515F: 5’-GTGYCAGCMGCCGCGGTAA-3’; 806R: 5’-GGACTACNVGGGTWTCTAAT-3’) in 30-cycle PCR reactions containing 15 μL Phusion High-Fidelity Master Mix (Thermo Scientific, USA) and 0.2 μM primers. PCR products underwent magnetic bead-based purification (AMPure XP, Beckman Coulter) and were normalized by fluorometric quantification (Qubit 4.0, Thermo Fisher). Libraries were prepared through end-repair, adapter ligation, and size selection (300-400 bp) using BluePippin (Sage Science). Library quality was verified through bioanalyzer electrophoresis before 2×150 bp paired-end sequencing on an Illumina NovaSeq 6000 platform. Bioinformatic processing involved demultiplexing raw reads using sample-specific barcodes, followed by read merging with FLASH (v1.2.11) and quality filtering via fastp (v0.23.1) to obtain high-quality clean tags. Chimeric sequences were removed using VSEARCH (v2.16.0) against SILVA reference databases. Amplicon Sequence Variants (ASVs) were resolved through DADA2 denoising in QIIME2, with taxonomic classification performed against SILVA database.

### Statistical analysis

Statistical analysis was performed using GraphPad Prism 10. Experimental results were displayed as mean ± SD or mean ± SEM. Differences between two groups were tested with an unpaired two-tailed Student’s t-test. Differences between survival curves were analyzed using a log-rank test.

## Acknowledgments

This study was supported by National Key R&D Program of China (2023YFA0915300 to X.Y. and 2023YFA0914900 to X.X.), National Natural Science Foundation of China (22025207 to X.Y., 22377127 to R.X. and 52473159 to T.L.), Strategy Priority Research Program (Category C) of Chinese Academy of Sciences (XDC0290200 to T.L.), Autonomous Deployment Project of State Key Laboratory of Biopharmaceutical Preparation and Delivery (2024-FX-A-01 to T.L.).

## Extended Data

**Extended Data Fig. 1.**
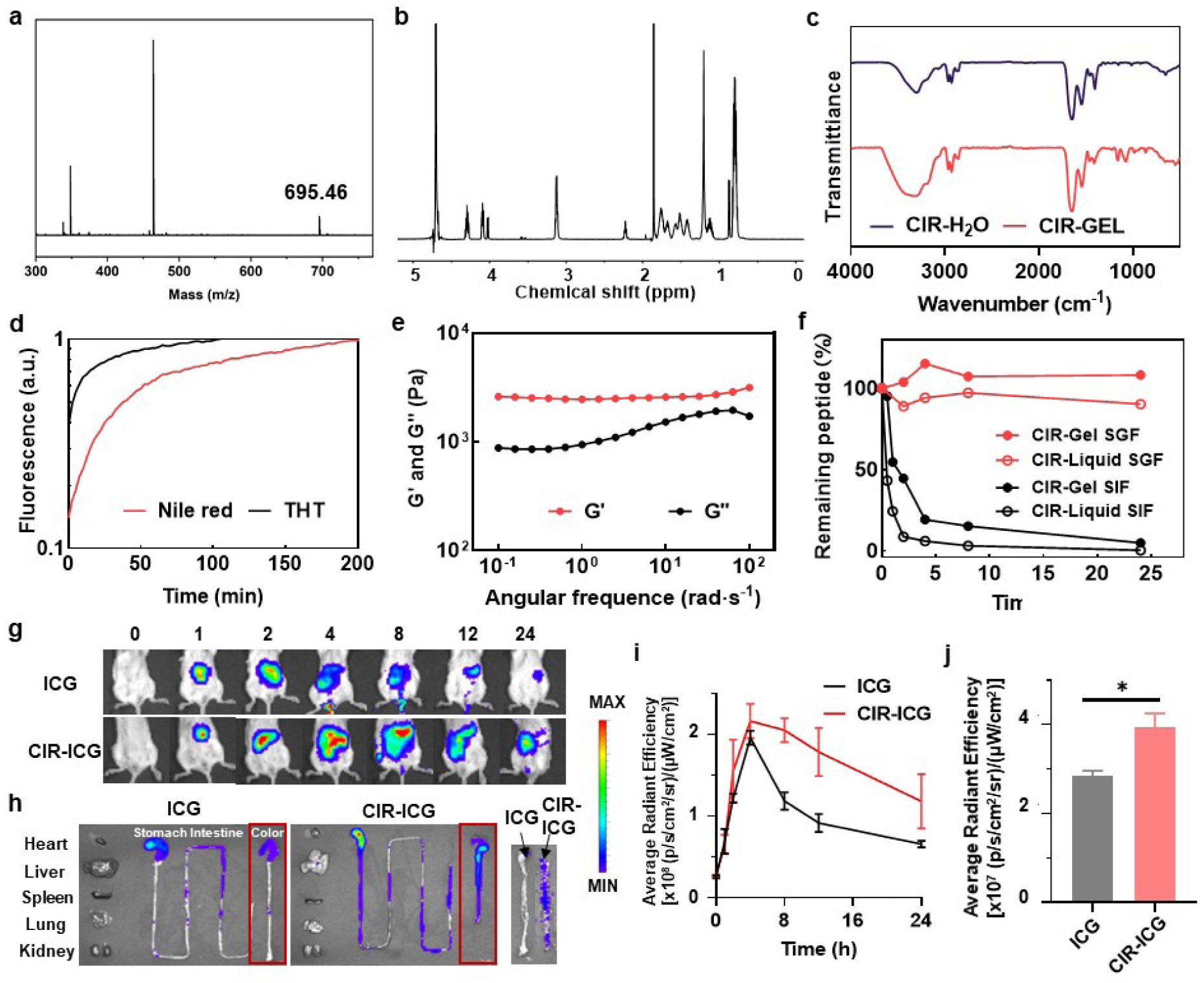
Physicochemical properties and oral-delivery stability of CIR hydrogel. **a**, Mass spectrum of CIR peptide. **b**, ^1^H-NMR spectrum of CIR peptide. **c**, FT-IR spectrum of CIR peptide and hydrogel. **d**, ThT and Nile red fluorescence labelling of CIR hydrogels with dynamic measurements showing that the gelation process was driven by the formation of hydrogen bonds and hydrophobic interactions. **e**, Rheological analysis of CIR hydrogels with frequency sweep demonstrating that CIR self-assembled into hydrogels with robust mechanical strength. **f**, Degradability of the CIR hydrogel and liquid in simulated gastric fluid (SGF) and simulated intestinal fluid (SIF), showing improved stability of CIR after gelation. **g**, In vivo imaging of mouse abdomen at different times after oral administration of ICG and CIR hydrogel encapsulating ICG. **h**, Ex vivo imaging of main organs of mouse at 24 h after oral administration of ICG and CIR hydrogel encapsulating ICG. Right: Colon tissues with feces removed. **i,j**, Quantified radiant efficiency of in vivo (i) and ex vivo (j) imaging of colon tissues, which demonstrated the improved retention of CIR hydrogel in colon. * p < 0.05, by Student’s t test.

**Extended Data Fig. 2.**
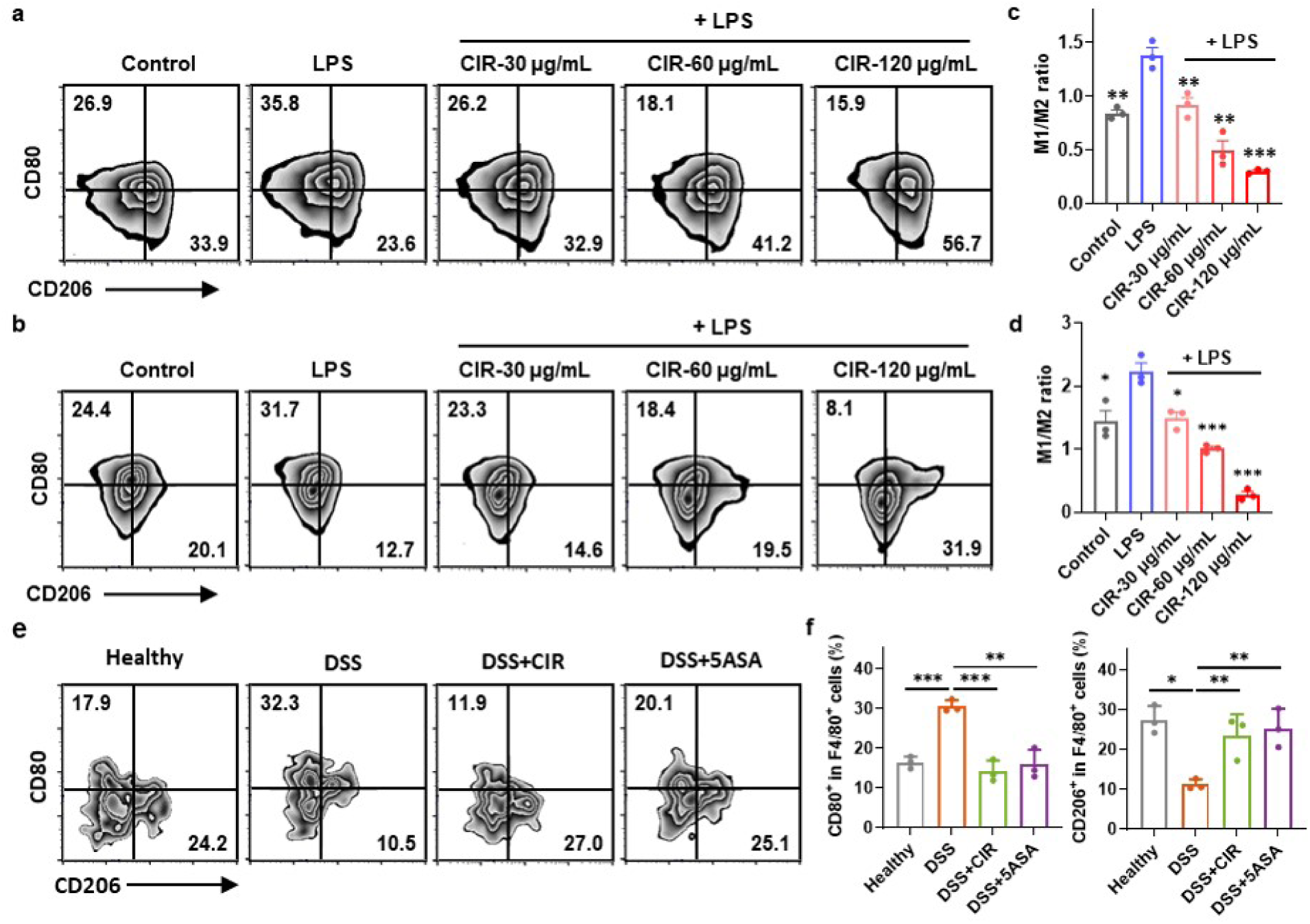
CIR hydrogel modulated macrophage differentiation and inhibited inflammatory cytokine release. **a,b,** Flow cytometry of BMDM (a) and RAW264.7 (b) under the treatment of LPS and CIR hydrogels in different concentrations showing the expression levels of CD80 and CD206. **c,d,** Quantification of macrophage subtype M1/M2 ratio from flow cytometry results of BMDM (c) and RAW264.7 (d) demonstrated that CIR hydrogel induced macrophage differentiation to M2 subtype. **e,** Flow cytometry of macrophages extracted from mouse spleens in acute colitis model under different treatments. **f,** Quantification of CD80^+^ and CD206^+^ cells in F4/80^+^ cells from flow cytometry results of mouse spleen tissues. Data represent mean ± SEM. * p < 0.05, ** p < 0.01, *** p < 0.001, by Student’s t test. Comparisons are with the LPS-only group if unlabeled.

**Extended Data Fig. 3.**
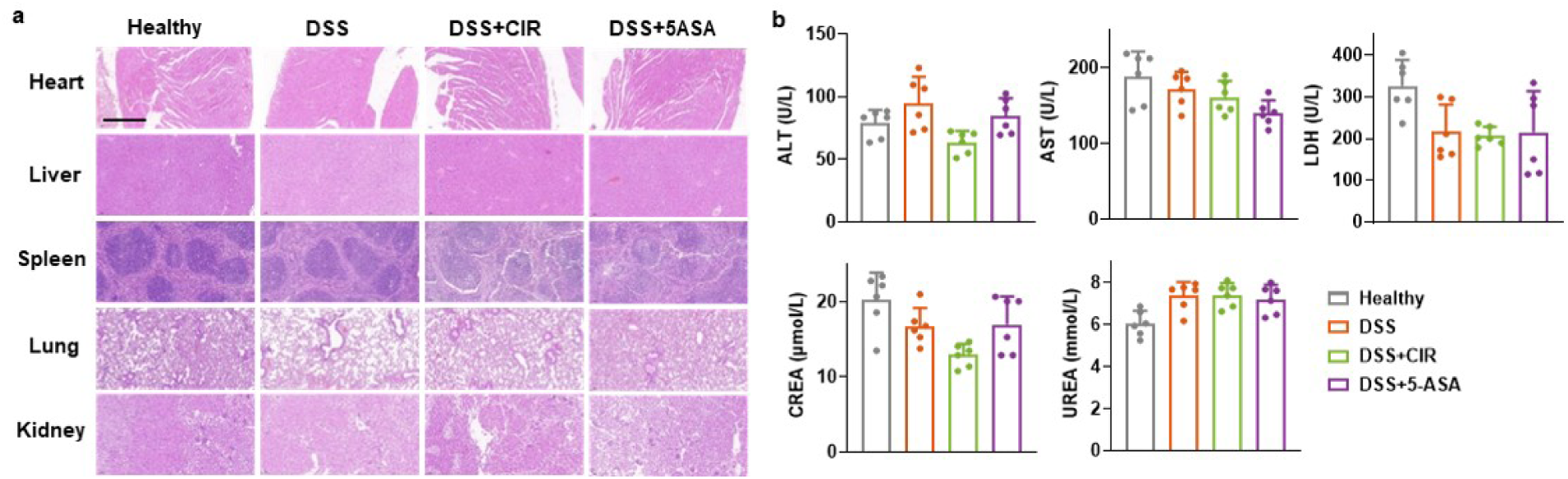
Biosafety evaluation of CIR hydrogels in animal models. **a,** Histopathological images of heart, liver, spleen, lung, and kidney in mouse model revealed no evident tissue damage attributable to CIR hydrogel treatment. Scale bar, 500 μm. **b,** Mouse serum biochemical tests showed no significant alterations in key serum markers, including alanine aminotransferase (ALT), aspartate aminotransferase (AST), lactate dehydrogenase (LDH), creatinine (CREA), and urea. Data represent means ± SEM. * p < 0.05, ** p < 0.01, *** p < 0.001, by Student’s t test.

**Extended Data Fig. 4.**
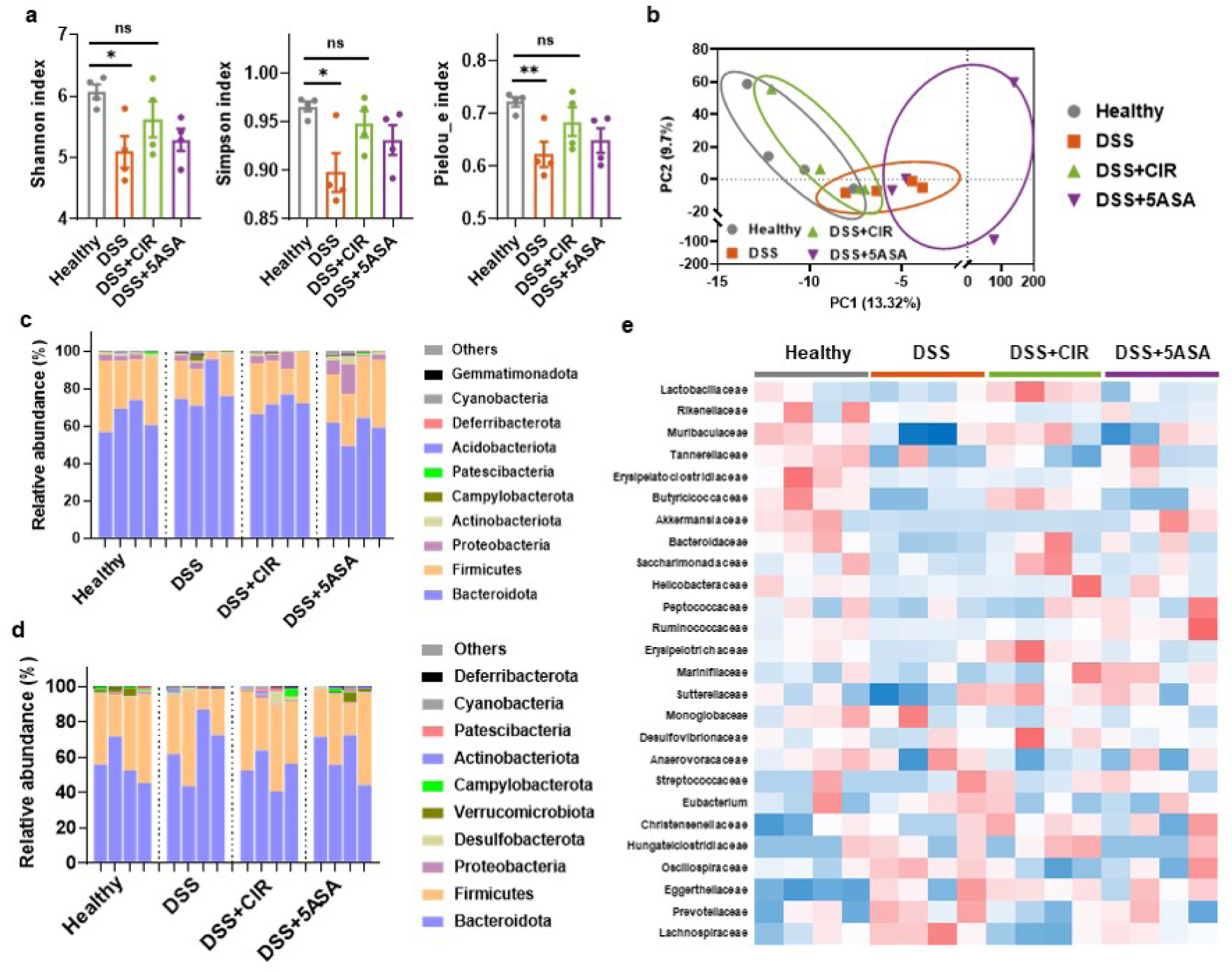
CIR hydrogels modulated microbiota abundance and diversity in UC mouse and NHP models. **a,** Venn diagram showing the number of unique and overlapped OTUs among Healthy, DSS, and DSS+CIR groups in the acute colitis mouse model. **b,** Radar diagram showing α diversity with Shannon, Simpson, Pielou_e, and Chao1 index in the acute colitis mouse model. **c,** Alpha diversity of microbiome in the chronic colitis mouse model, including Shannon, Simpson, and Pielou_e index. **d,e,** PCA figures of microbiome in the acute (d) and chronic (e) colitis mouse model showing the *β*-diversity, each point represents one mouse (n = 4). **f-h,** Relative abundance of microbiome that is significantly altered at phylum levels in acute colitis mouse (f), chronic colitis mouse (g) and chronic colitis monkey (h) models. **i,j,** Heatmaps showing the microbiome abundance changes among different groups in acute (i) and chronic (j) colitis mouse models. Data represent means ± SEM. * p < 0.05, ** p < 0.01, *** p < 0.001, by Student’s t test.

**Extended Data Fig. 5.**
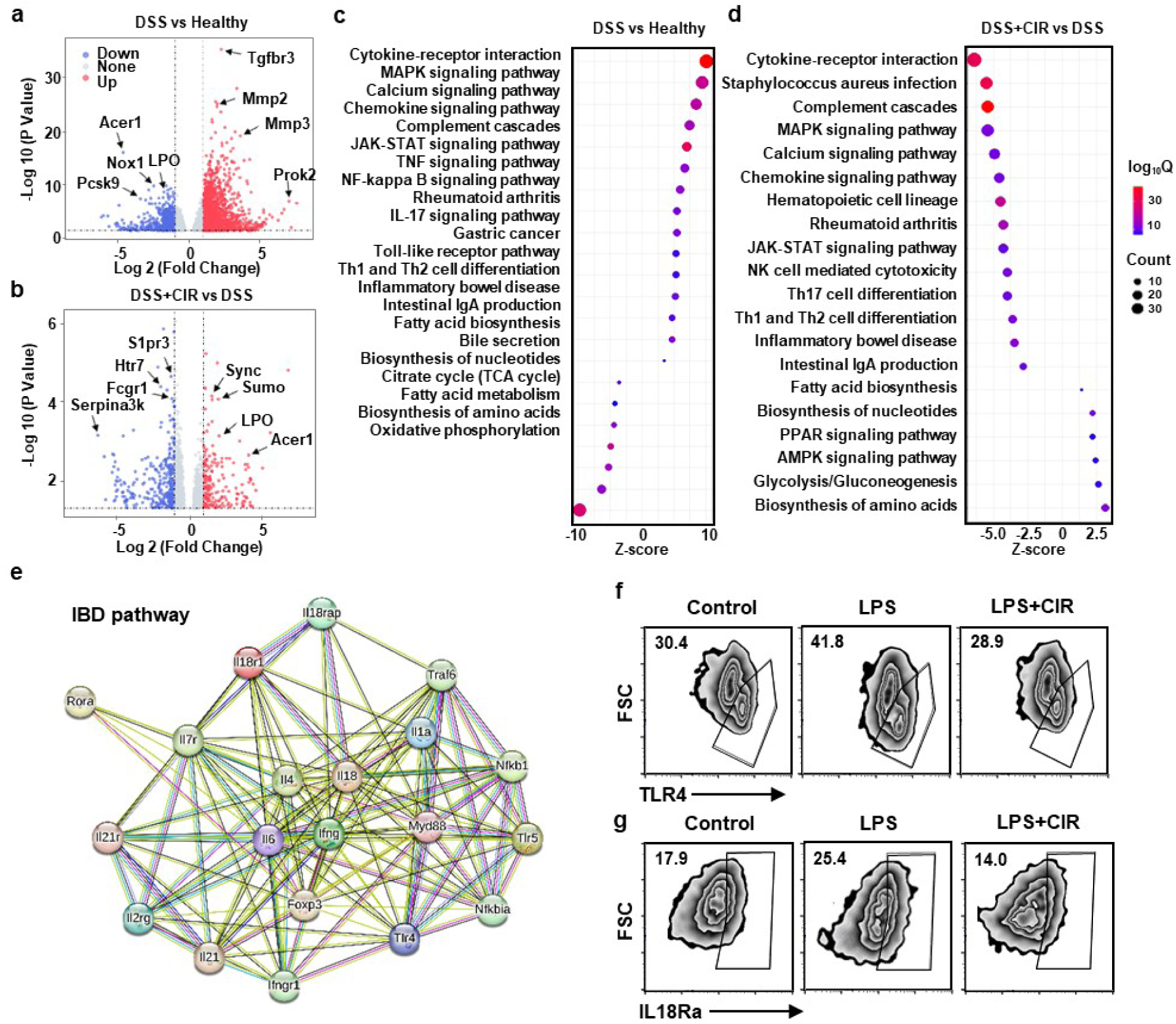
CIR hydrogels mitigated abnormal inflammation and immune response by downregulating TLR4 related pathways. **a,b,** RNA-seq of colon tissues from mice treated with DSS and DSS+CIR. Volcano plot for DEGs between DSS and Healthy groups (a) and between DSS+CIR and DSS groups (b). **c,d,** KEGG enrichment analysis revealing the altered inflammatory and immune signaling pathways in DSS (c) and DSS+CIR (d) groups. **e,** PPI analysis of DEGs in IBD pathway exhibiting interactions among the highly differentiated genes. **f,g,** Flow cytometry showing the expression levels of TLR4 (f) and IL-18Ra (g) in BMDM under the treatment of LPS and CIR peptide.

